## Supplementary Materials for "Sulcal morphology of posteromedial cortex substantially differs between humans and chimpanzees"

Willbrand, Maboudian *et al.*

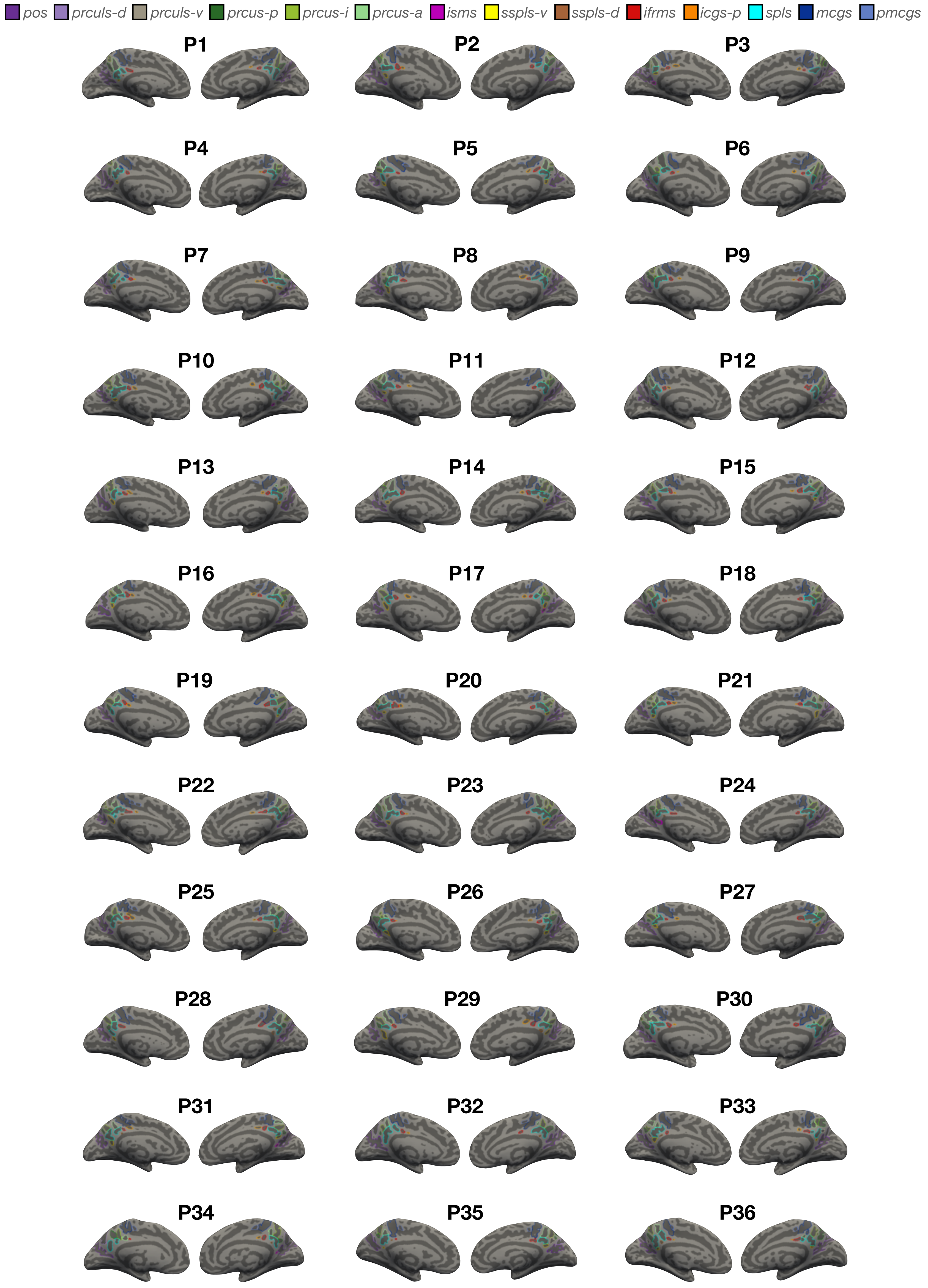
pos
prculs-d
prculs-v
prcus-p
prcus-i
prcus-a
isms
sspls-v
sspls-d
ifrms
icgs-p
spls
mcgs
pmcgs
P1
P2
P3
P4
P5
P6
P7
P8
P9
P10
P11
P12
P13
P14
P15
P16
P17
P18
P19
P20
P21
P22
P23
P24
P25
P26
P27
P28
P29
P30
P31
P32
P33
P34
P35
P36

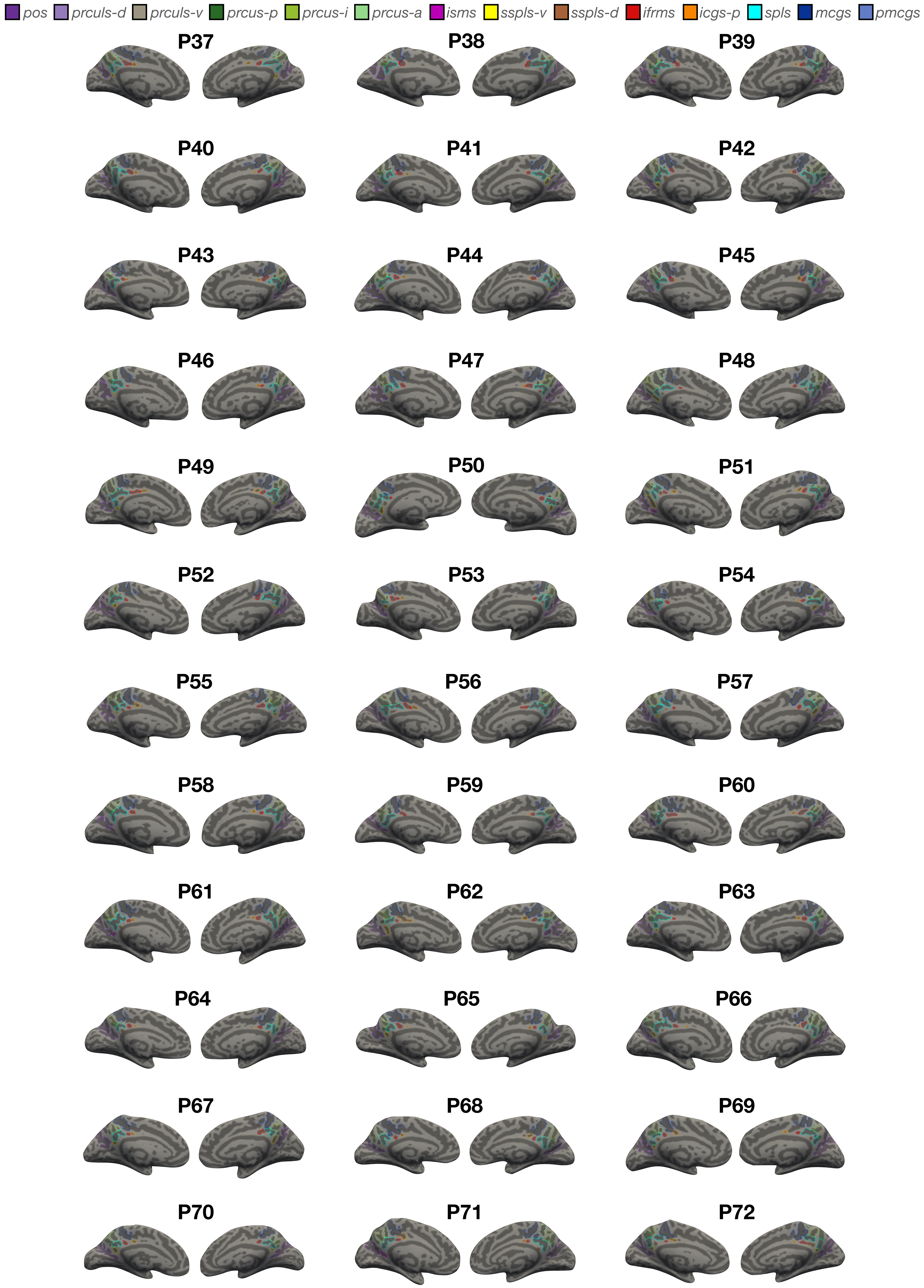


**Supplementary Figure 1. Manual PMC sulcal labels in every human participant.** Each sulcus is displayed on the left and right hemisphere inflated cortical surfaces in FreeSurfer 6.0.0, with label displayed as an outline according to the key at the top. Each hemisphere contains at least 8 sulci (from posterior to anterior): pos, prculs-d, prcus-p, prcus-i, prcus-a, spls, mcgs, and ifrms. An additional 6 sulci are variably present: isms, prculs-v, sspls-v, sspls-d, icgs-p, and pmcgs.

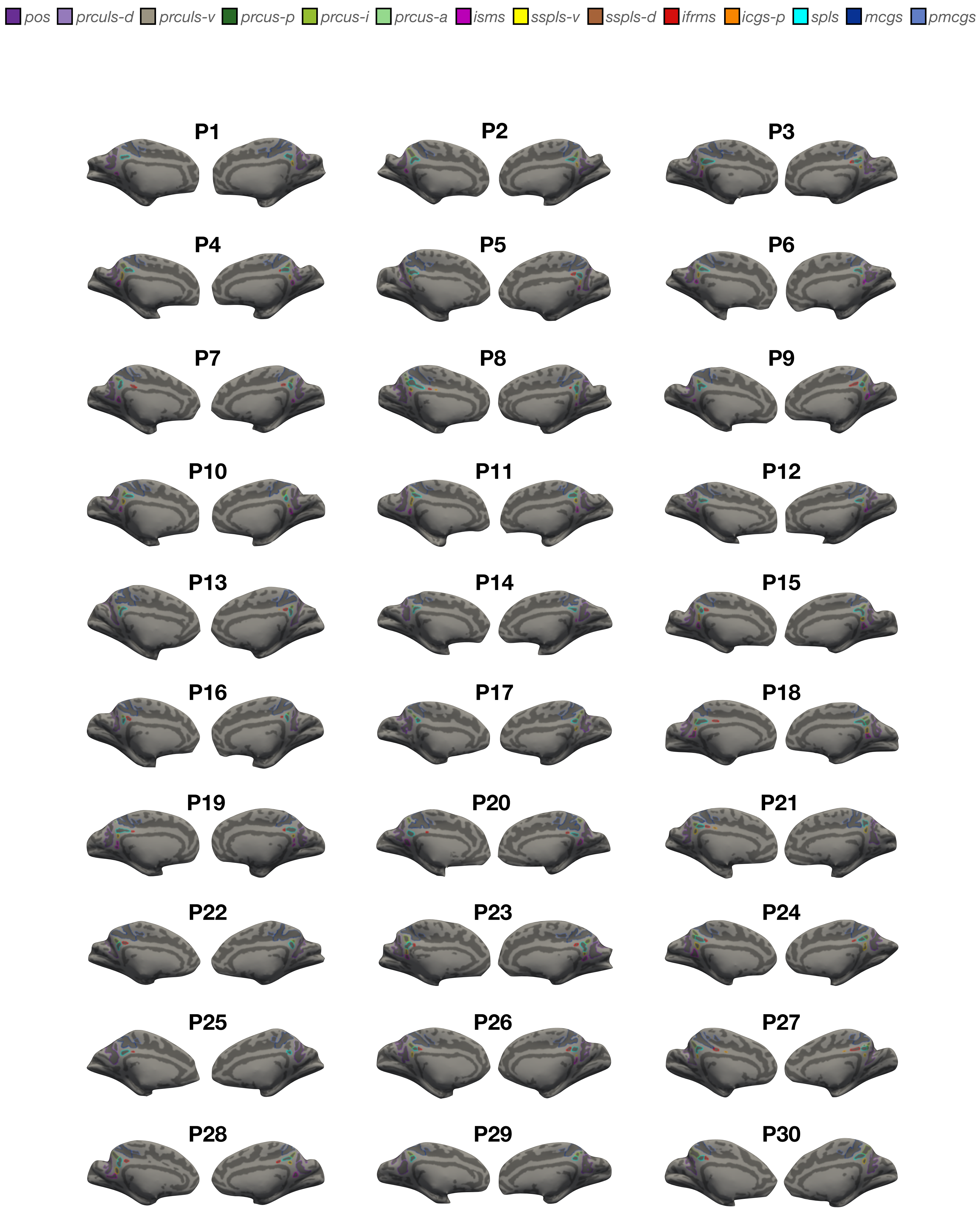
pos
prculs-d
prculs-v
prcus-p
prcus-i
prcus-a
isms
sspls-v
sspls-d
ifrms
icgs-p
spls
mcgs
pmcgs
P1
P2
P3
P4
P5
P6
P7
P8
P9
P10
P11
P12
P13
P14
P15
P16
P17
P18
P19
P20
P21
P22
P23
P24
P25
P26
P27
P28
P29
P30

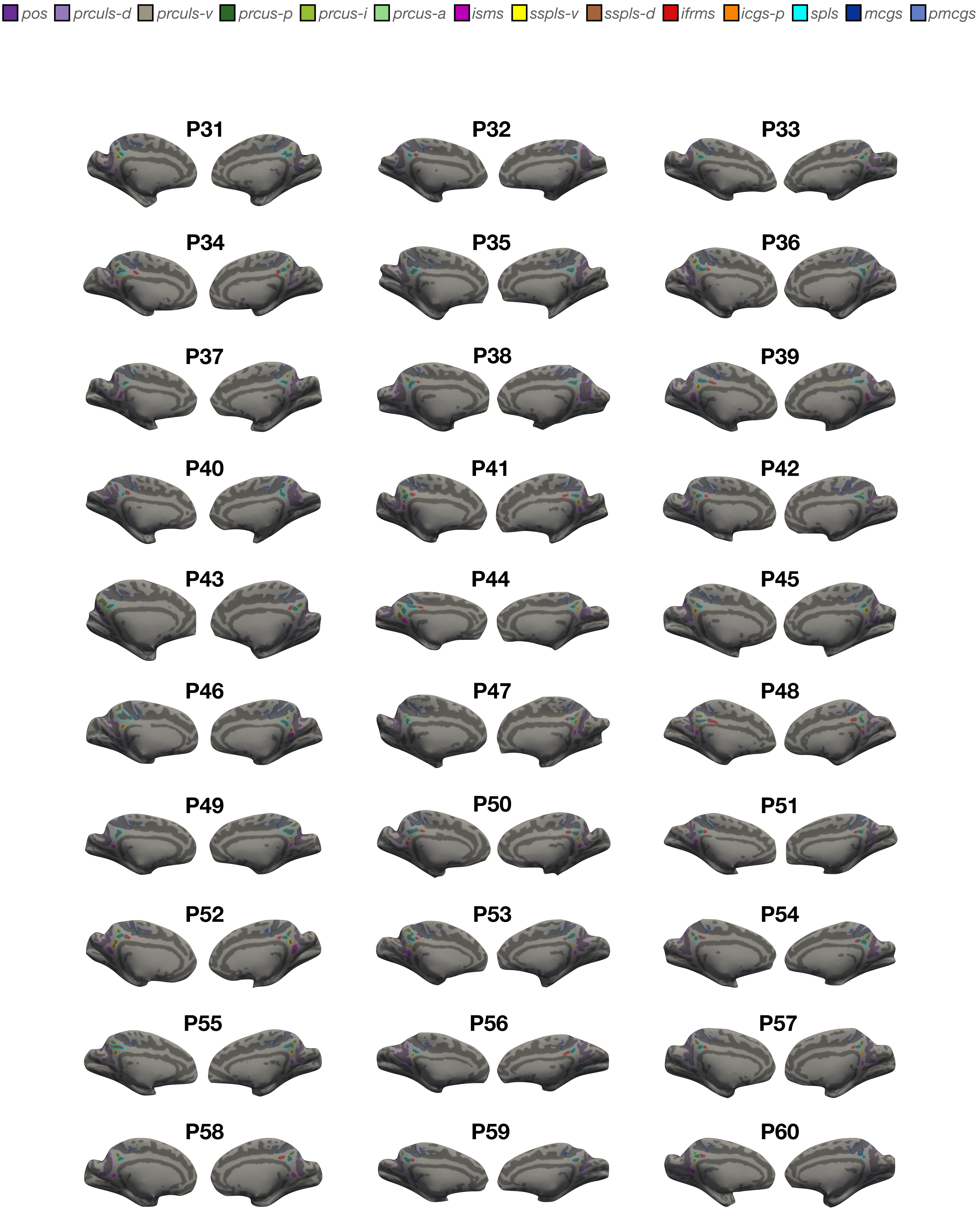


**Supplementary Figure 2. Manual PMC sulcal labels in every chimpanzee.** Each sulcus is displayed on the left and right hemisphere inflated cortical surfaces in FreeSurfer 6.0.0, with label displayed as an outline according to the key at the top. Each hemisphere contains at least 4 sulci (from posterior to anterior): mcgs, pmcgs, spls, and pos. An additional 8 sulci are variably present: prculs-d, prculs-p, prcus-i, prcus-a, isms, sspls-v, ifrms, and icgs-p. (2 sulci present in humans are not present in any chimp hemispheres: prculs-v, sspls-d).

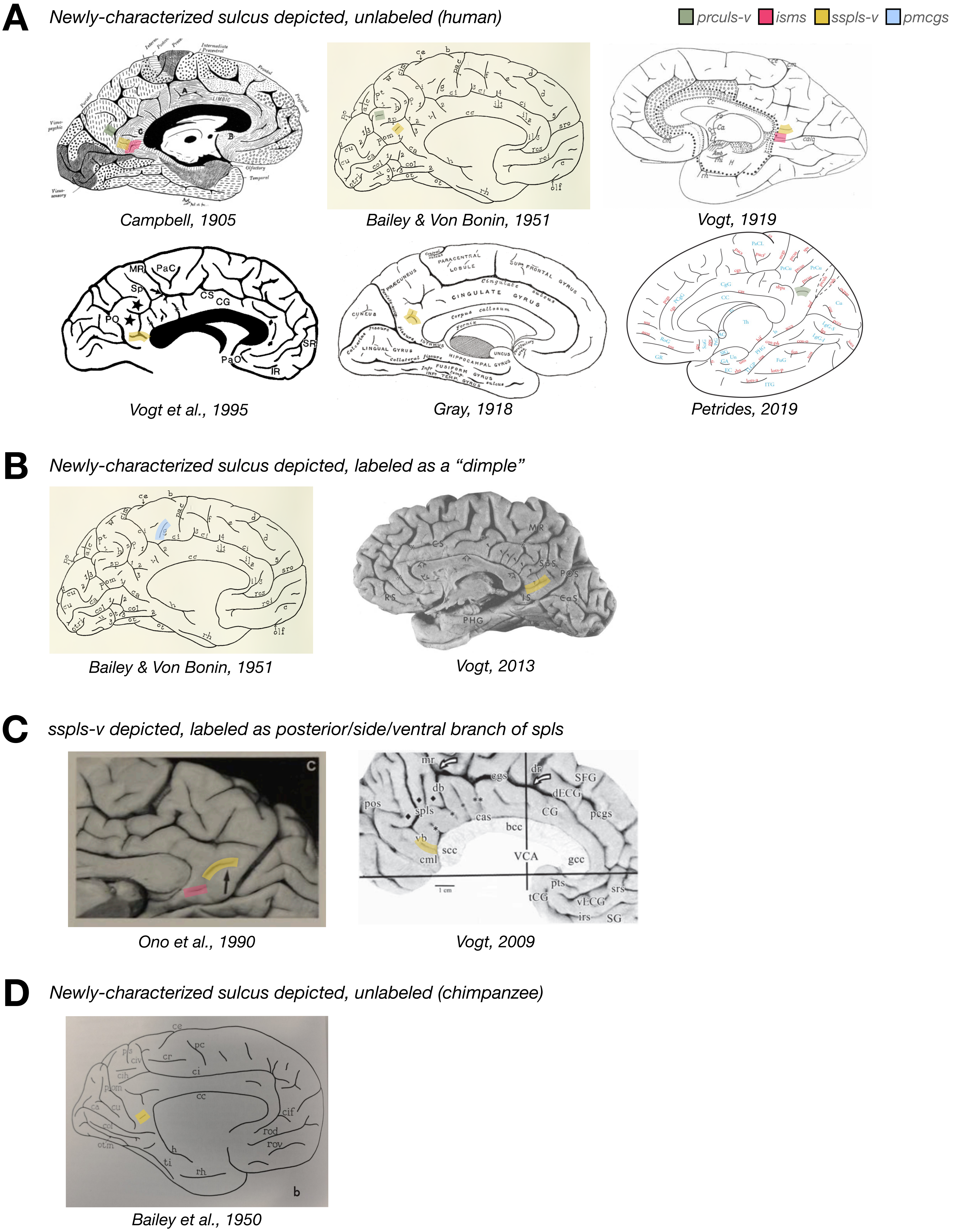


**Supplementary Figure 3. Previous depictions of newly-characterized indentations by anatomists.** While we label and quantify the incidence rates of four sulci (prculs-v, isms, sspls-v, and pmcgs) across species for the first time, some classical and modern anatomists have included an unlabeled (**A**) sulcus or (**B**) dimple in the location of some of these sulci in their schematics of human brains. **C.** In some modern studies, sspls-v has been labeled as part of the ventral or posterior branch of spls. **D.** Sspls-v has also been depicted unlabeled in chimpanzee brains. In all figures, colored shading has been added to show the sulcal label used for each indentation in the present study.

*pos* *prculs-d* *prculs-v* *prcus-p* *prcus-i* *prcus-a* *isms* *sspls-v* *sspls-d* *ifrms* *icgs-p* *spls* *mcgs* *pmcgs*

*Nbr 3* | *Nbr 3*

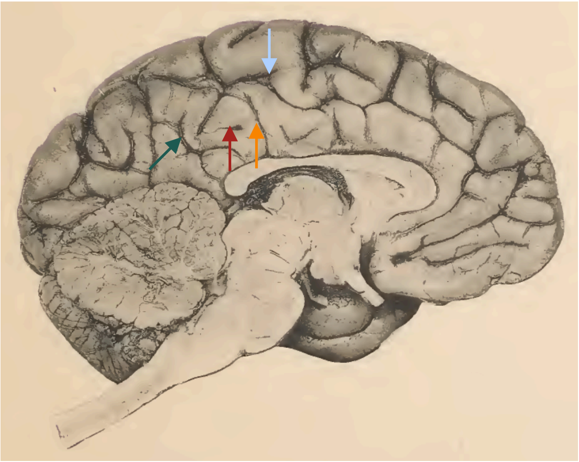

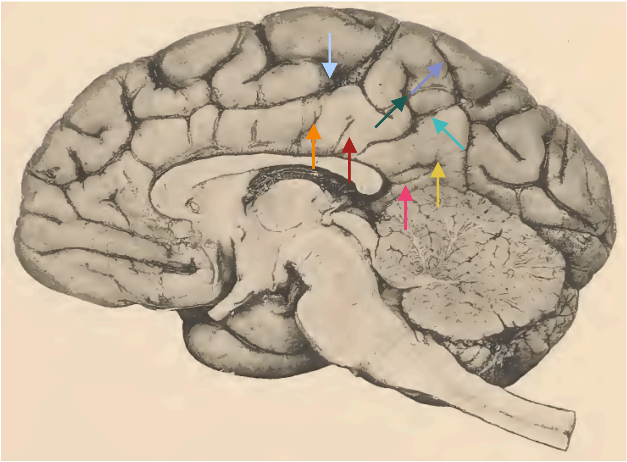

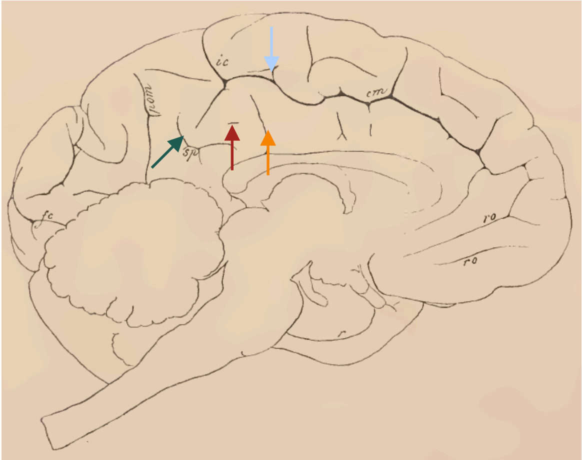


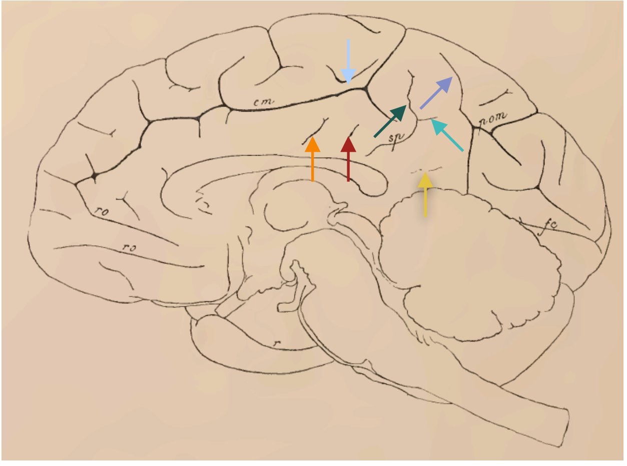


*Nbr 4* | *Nbr 4*

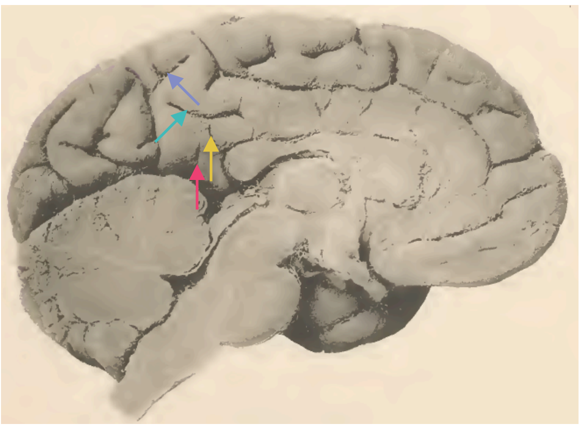

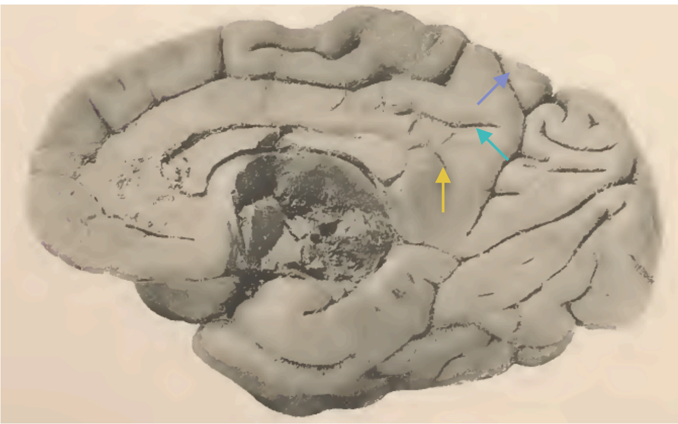

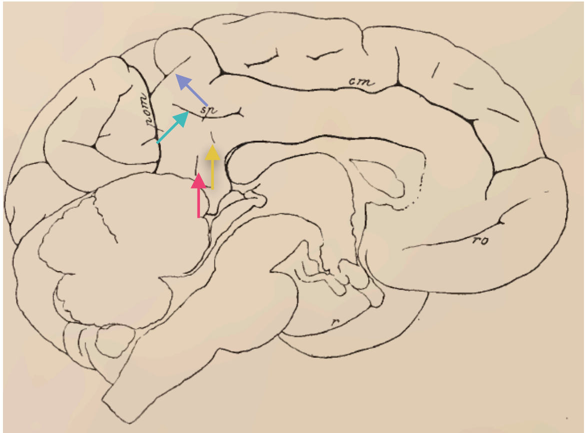


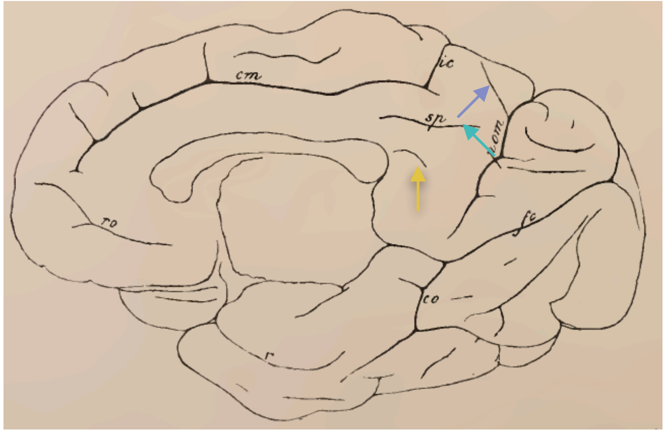

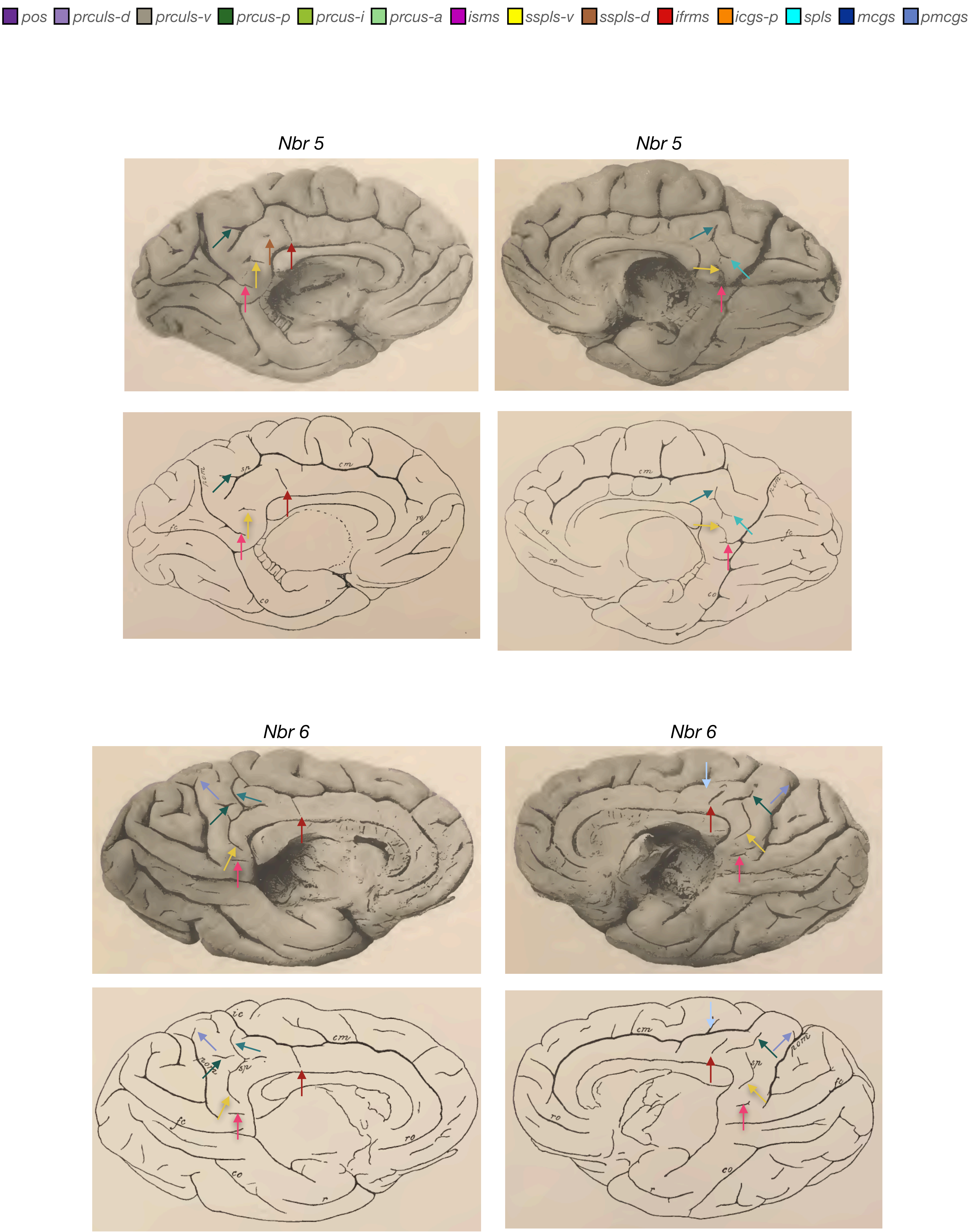
pos
prculs-d
prculs-v
prcus-p
prcus-i
prcus-a
isms
sspls-v
sspls-d
ifrms
icgs-p
spls
mcgs
pmcgs
Nbr 5
Nbr 5
Nbr 6
Nbr 6
pom
sp
cm
fc
ro
co
ic

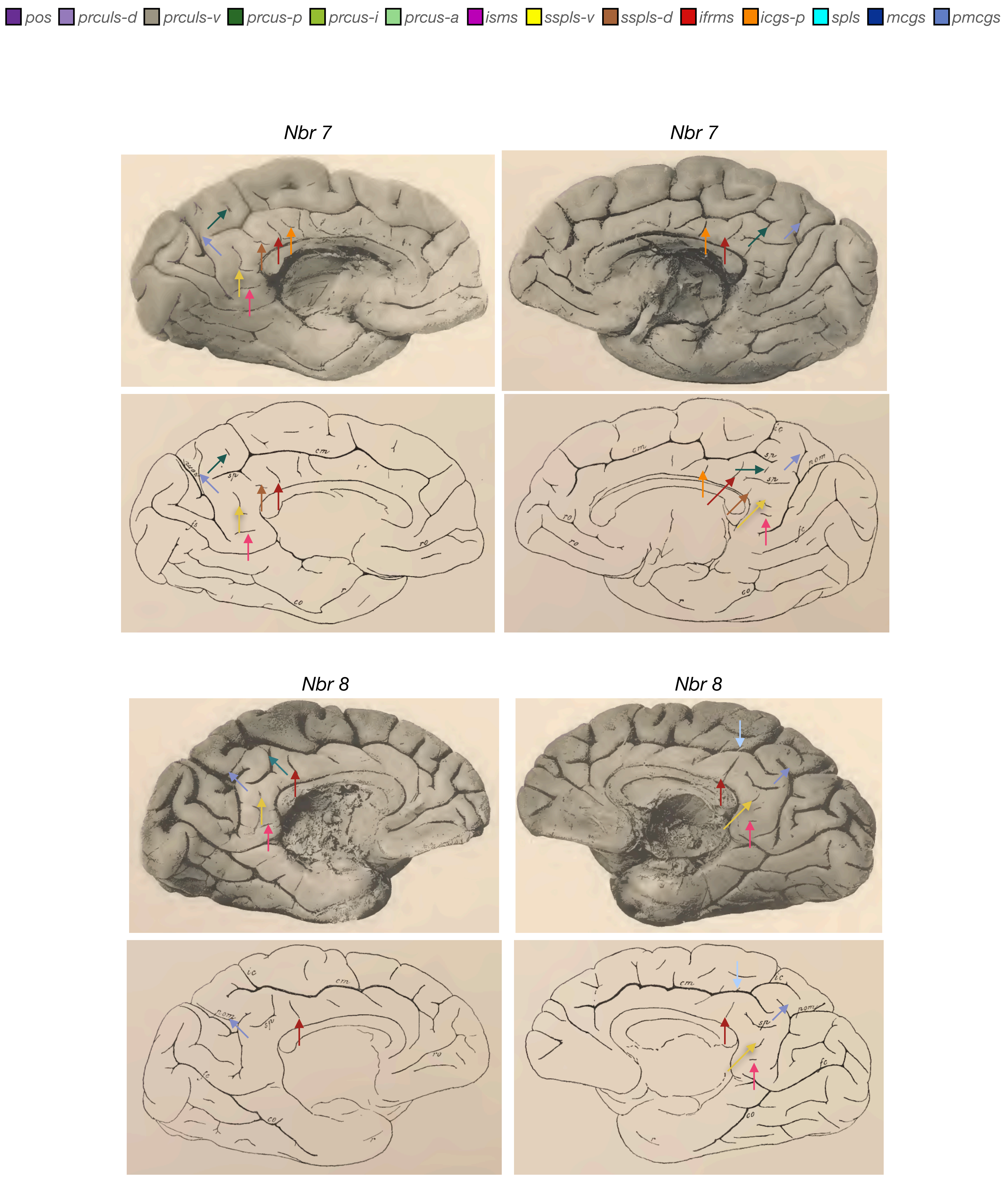

pos
prculs-d
prculs-v
prcus-p
prcus-i
prcus-a
isms
sspls-v
sspls-d
ifrms
icgs-p
spls
mcgs
pmcgs
Nbr 7
Nbr 7
Nbr 8
Nbr 8

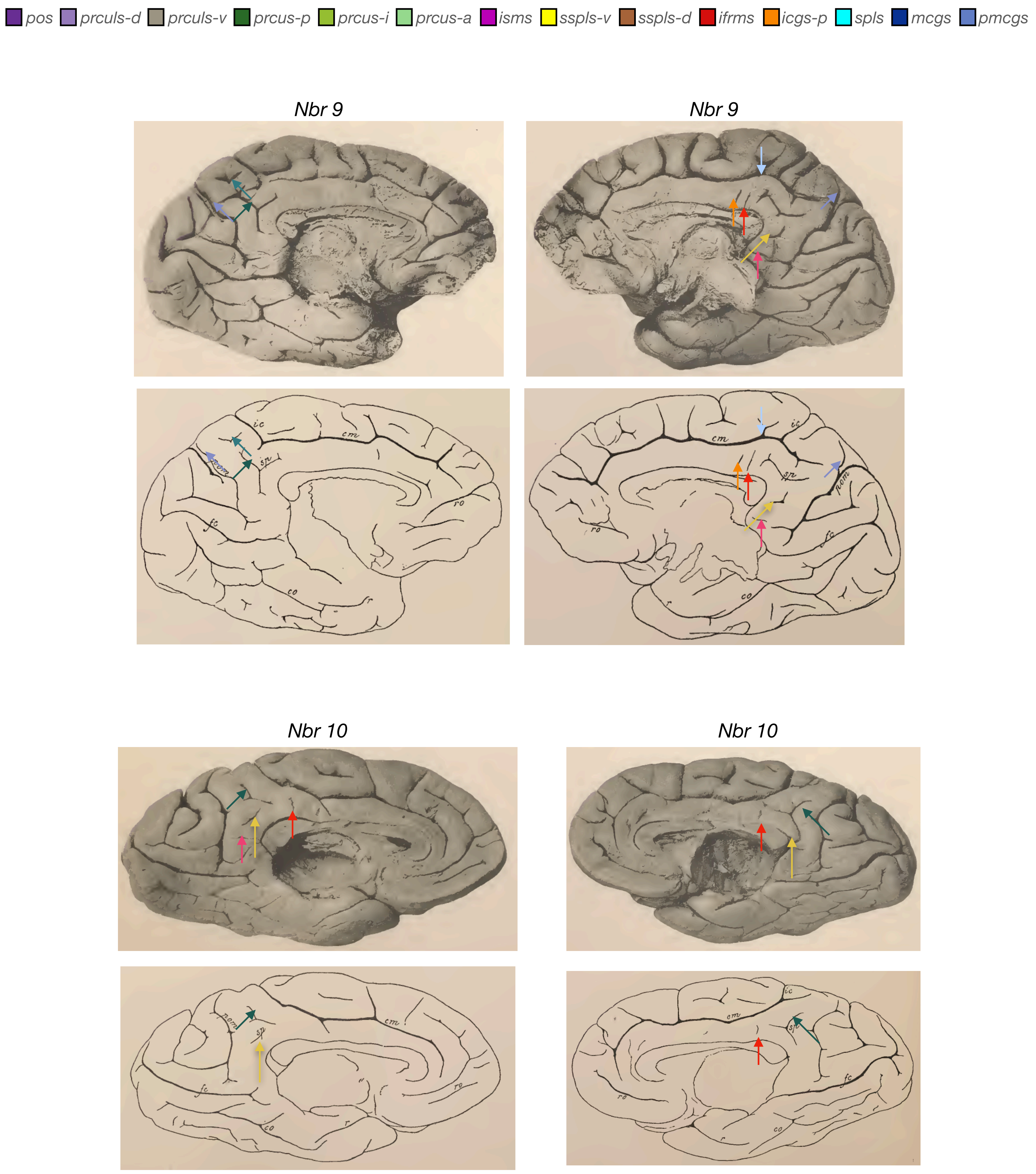
pos
prculs-d
prculs-v
prcus-p
prcus-i
prcus-a
isms
sspls-v
sspls-d
ifrms
icgs-p
spls
mcgs
pmcgs
Nbr 9
Nbr 9
Nbr 10
Nbr 10

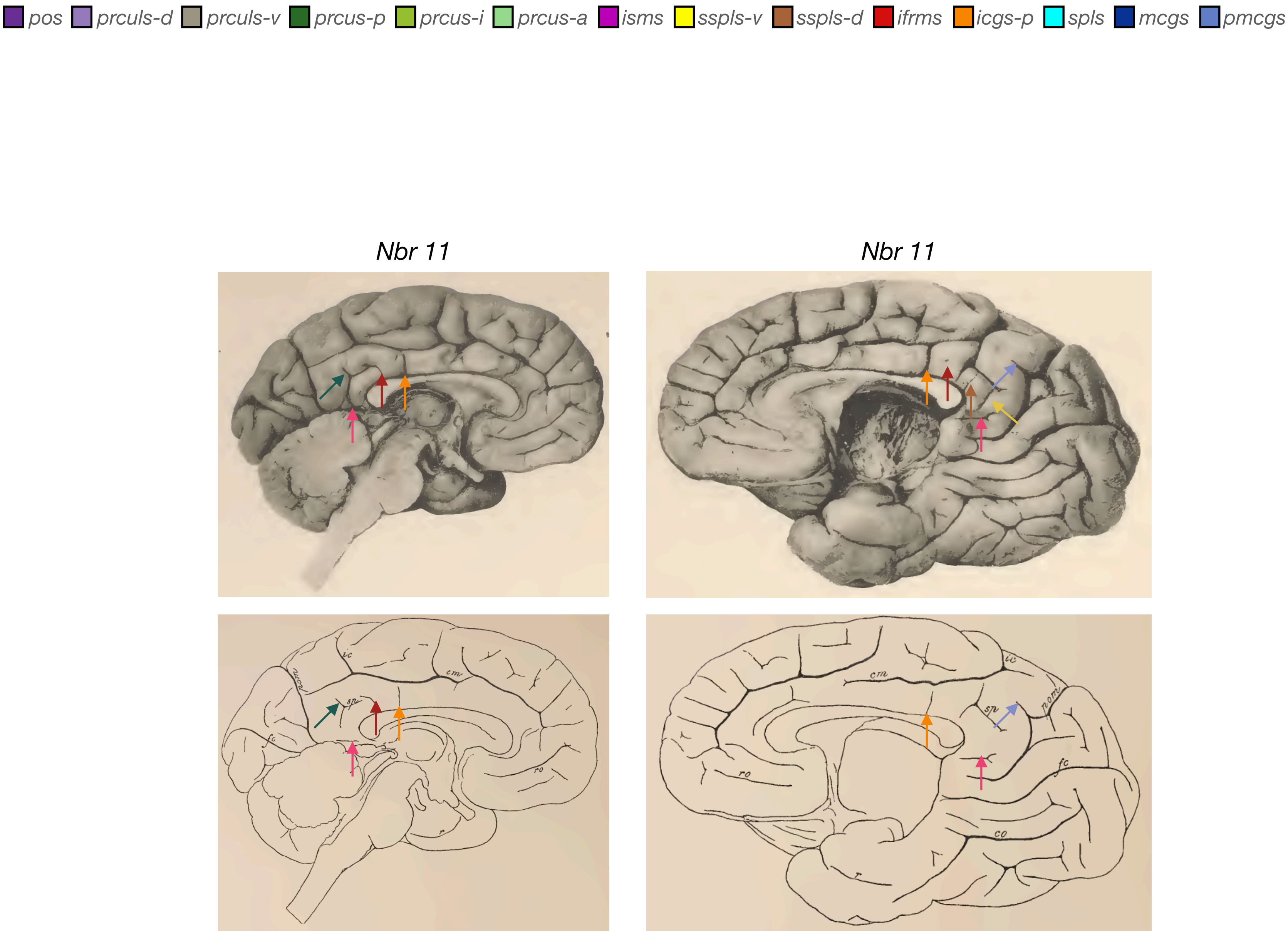


**Supplementary Figure 4. Manual PMC sulcal labels in the left and right hemispheres of postmortem chimpanzee brains.** Nine postmortem brains and their respective schematics from Retzius' 1906 atlas. PMC sulci are defined with arrows colored according to the key at the top of the figure. We do not mark the pos, spls, and mcgs.
